# Assessment of antimicrobial resistance genes in Caribbean corals, including those treated with amoxicillin

**DOI:** 10.1101/2024.11.01.621504

**Authors:** Karen L. Neely, Christina A. Kellogg, Julie J. Voelschow, Allison R. Cauvin, Sydney A. M. Reed, Ewelina Rubin, Julie L. Meyer

## Abstract

The decimation of reefs caused by stony coral tissue loss disease prompted the use of a topical amoxicillin treatment to prevent coral mortality. Application of this treatment led to concerns about unintentional impacts such as potential alteration of the coral microbiome and possible spread of antibiotic resistance. We used two different methodologies – microbial RNA sequencing and microbial qPCR array – to assess these concerns and to establish a baseline of antimicrobial resistance genes in coral microbes. RNA sequencing was conducted on coral mucus samples collected before and 24 hours after amoxicillin application on wild *Montastraea cavernosa*. While diverse antibiotic resistance genes (ARGs) were expressed, no differences were detected in ARG expression after amoxicillin treatment. Additionally, there were no notable changes in the microbial communities between the before and after samples. Microbial qPCR array was used to assess differences in ARGs over longer timescales in wild *Colpophyllia natans*, comparing never-treated corals with ones treated a single time seven months prior and with those treated multiple times seven months and more prior. No clinically relevant ARGs represented in the arrays were detected across any samples. A small number of above-detection reads (4 in the never-treated corals, 2 in the once-treated corals, and 0 in the multi-treated corals) may indicate weak amplification of similar environmental (non-anthropogenic) ARGs in the corals. Results indicate that the localized topical application of amoxicillin to prevent mortality of SCTLD-affected corals is neither immediately disrupting the microbiome of treated corals at the colony level nor driving the development of ARGs over immediate or longer-term time scales.

## 1. Introduction

The outbreak of stony coral tissue loss disease (SCTLD) has caused unprecedented mortality across the majority of stony coral species in the Caribbean, including more than 20 of the approximately 45 species of reef-builders and 5 of the U.S. Endangered Species-listed Caribbean coral species (Florida Coral Disease Response Research & Epidemiology Team 2018). The disease originated in South Florida in 2014 (Precht et al. 2016), but since 2017 has spread throughout much of the greater Caribbean region (Papke et al. 2024).

Mortality rates of 67-100% of some species (Precht et al. 2016), along with significant losses to coral cover, diversity, and ecosystem function (Alvarez-Filip et al. 2022, Hayes et al. 2022), triggered unprecedented efforts to save individual colonies using novel methods. Antibiotics added to an aquarium with SCTLD-affected corals halted disease progression (Miller et al. 2020), and a topical paste was subsequently developed specifically for delivering amoxicillin to diseased coral tissues. The paste, termed amoxicillin+Base2b, was tested in the field on various species and found to be 67–100% effective at halting SCTLD lesions (Neely et al. 2020, Neely et al. 2021, Shilling et al. 2021, Walker et al. 2021). Though in some instances new lesions can appear on previously treated corals, sites that are regularly visited can address these, leading to high survival of corals (Neely 2023). The application of this antibiotic paste has since become the standard for treatment of SCTLD throughout the Caribbean. As of May 2025, over 30,000 corals have been treated with this method on Florida’s Coral Reef (Florida Fish and Wildlife Research Institute 2024).

Though these successes are encouraging, there are management concerns about the deployment of antibiotics onto coral reefs. These include the short- and/or long-term disruption of the coral microbiome, as well as the potential short- and/or long-term development of antibiotic resistance within the corals and/or the surrounding environment. Assessments of these potential unintended consequences were identified by the four leading state and federal agencies as research priorities for Florida’s Coral Disease Response in 2021, 2022, and 2023. The concerns also delayed or prohibited use of these treatments in some Caribbean jurisdictions (Lee Hing et al. 2022).

The widespread usage of antibiotics in medical and agricultural fields has prompted great concerns over the development of antibiotic resistance in our environment (Kümmerer 2003, Zhang et al. 2022). While the appearance of resistant microbes over the course of more than 80 years of human use has doubtless been exacerbated by the development and use of antibiotics, studies have also revealed that antibiotic resistance is ancient (Waglechner et al. 2021), detecting a variety of antibiotic resistant genes (ARGs, also known as AMRs) in 30,000-year-old permafrost and deep within caves that have been isolated for millions of years (D’Costa et al.

2011, Bhullar et al. 2012). The likely ubiquitous presence of ARGs in natural systems relates to their extensive role in common competitive microbe-microbe interactions (Granato et al. 2019) as well as additional physiological functions outside of their protective role (e.g., encoding efflux pumps or signal trafficking; Martínez (2008)). Nevertheless, anthropogenic antibiotic use strongly influences microbial genetic selection which can result in clinically relevant antibiotic resistance.

Antibiotics are commonly used in the treatment of human ailments as well as illnesses of pets and livestock. Additionally, they are frequently used prophylactically in agriculture and aquaculture or as growth promoters (Kümmerer 2003, Cabello 2006, Kümmerer 2009a).

Estimates from the late 1990s and early 2000s placed worldwide consumption of antibiotics at 100,000 to 200,000 tons per year, with as little as 10% being for human clinical use (Kümmerer 2009b). Due to concerns that overuse of antibiotics in animal feeds could compromise their use in human medicine, their use as growth promoters has been banned in a number of European countries (Kümmerer 2009b). Given that a bacterium under optimal conditions can divide as rapidly as every 20 to 30 min, it only takes about 20 generations to reach over 1 million cells; so theoretically a cell containing an ARG could become dominant in a population within as little as 10 hours (Pray 2008). Use of antibiotics for treatment of wildlife diseases, however, is uncommon. In general, disease outbreaks in free-ranging wildlife are considered natural and left to run their course. Exceptions include when those outbreaks may also affect human, pet, or agricultural health, or are impacting highly endangered or managed species (as reviewed in Neely et al. (2021)).

It is currently unknown what sort of selective effects the use of topical amoxicillin may have on coral microbiomes or ARGs (Saldaña and Woodley 2024), nor what kind of background levels and diversity of these genes might already exist in corals. Here we tested whether the application of amoxicillin+Base2b on coral disease lesions caused a change in the coral microbiome in tissues adjacent to the lesion 24 hours after application. We also conducted two different experiments using two different methods – metatranscriptomics and quantitative polymerase chain reaction (qPCR) microarrays – to assess whether there was an increase in the abundance or diversity of ARGs as a result of these small, periodic treatments. These experiments provide baseline information on the *in situ* presence and diversity of ARGs associated with corals in Florida and can inform future investigations of antibiotic applications in the marine environment.

## 2. Methods

### 2.1. Sample Sites and Collections

Coral tissue and mucus samples were taken from two patch reefs within the Upper Keys section of the Florida Keys National Marine Sanctuary: Hen and Chickens reef, and Cheeca Rocks. The two reefs are 7.5 km from each other (Figure 1a-c). Both lie 2-3 km from the nearest point of land, range in depth from 1.5 – 6.0 meters, and contain relatively high coral cover and coral species diversity compared to other Florida reefs. Here we compare the results of two independently designed and executed experiments, one from Hen and Chickens reef and one from Cheeca Rocks, to assess the in situ selective pressure for ARGs during antibiotic treatments for SCTLD.

**Figure 1.**
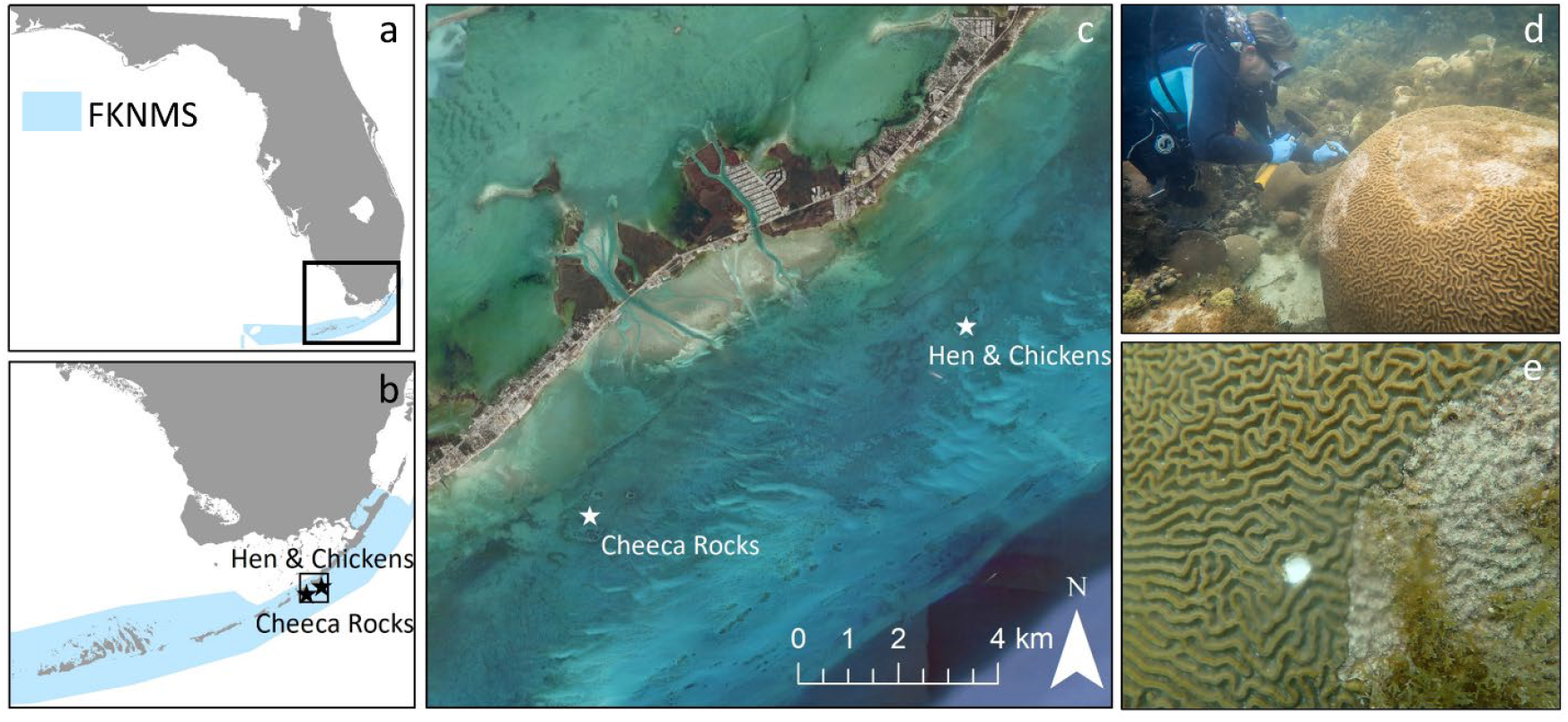
Map of sampling sites within the Florida Keys National Marine Sanctuary (a-c). Cores for microbial qPCR array were taken using a hole-punch (d) to remove a small area of tissue near previously treated stony coral tissue loss disease lesions (e).

At Hen and Chickens reef, sampling was designed to assess the baseline levels of ARGs and whether there were immediate changes in microbial community structure or in the expression of ARGs in coral tissues treated with antibiotics. Ten SCTLD-affected colonies of *Montastraea cavernosa*, all of which had never previously been observed with SCTLD and which had never been treated, were sampled on July 19, 2022. Mucus-tissue slurry samples were collected by aspiration with a sterile needle-less 20-ml syringe using gentle agitation of the surface mucus layer from apparently healthy tissue at least 30 cm from active SCTLD lesions. The surface mucus layer was chosen as the sample type since it is the interface between the coral holobiont and the environment and contains the highest density of prokaryotic cells (Garren and Azam 2010), thus providing the most active microbial community for detection of short-term changes in microbiome composition and gene expression. The sample was collected away from the active lesion in order to have a baseline reference for the coral microbiome that was unaffected by the disease lesion community. After sampling, any SCTLD lesions on the coral were treated with the standard amoxicillin+Base2b mixture (1:8 ratio by weight). The compound was topically applied directly to active disease lesions, overlapping the live-dead tissue margin and adhering to the skeleton and tissue. Twenty-four hours later, the same corals were revisited and sampled again, at a distance of three coral polyps from the treatment margin. The distance of three coral polyps (approximately 2 cm) was chosen as a representative distance for impacts to the coral microbiome directly outside of the treatment area. This sampling time frame was chosen because the treatment was designed to deliver a quick burst of antibiotic on the first day, followed by slower release over the next two days. Lab experiments show that more than half of the amoxicillin in the Base2b paste is delivered within 24 hours and all of the amoxicillin is delivered within 72 hours of application (Favero and Curtis 2020). In addition, 24 hours was hypothesized to be sufficient time for changes to be detectable in both microbiome composition and gene expression but not so long that the bacterial community recovered; Sweet et al. (2011) found orders of magnitude reduction in the bacterial population of corals following antibiotic water dosing, but substantial recovery between 48 and 96 hours. For both the before and after-treatment samples, syringes were immediately stored in a highly insulated cooler with dry ice. While this method keeps the samples very cold, the saltwater typically does not freeze. On shore, syringes with mucus-tissue slurries were set upright inside the cooler to allow settlement from concurrently sampled seawater. Excess seawater was decanted from the syringes, and the remaining mucus-tissue slurries were transferred to sterile conical tubes with Zymo DNA/RNA Shield. Samples were transferred frozen to the University of Florida and stored at −80°C until processing.

In a separate experiment at Cheeca Rocks, sampling was designed to assess whether antibiotic treatments were creating long-term selective pressure that would result in detectable antibiotic resistance genes, and whether single versus multiple treatments through time would affect this. Samples were taken from corals at Cheeca Rocks on November 17, 2022. The experimental design included sampling four colonies (biological replicates) within each of three categories: (i) never treated with amoxicillin+Base2b, (ii) treated a single time seven months prior to sampling, and (iii) treated at three sample times, at least seven months prior. Within humans, the rise of resistant pathogens can be delayed from 0 to 6 months after antibiotic treatments (Poku et al. 2023), and so the rationale for sampling several months after treatment in corals was to assess whether lasting resistance developed, particularly in the case of corals that had been treated multiple times.

A total of 12 samples from *Colpophyllia natans* were taken. Samples were also taken from 12 *Montastraea cavernosa* colonies following the same procedures, but the *M. cavernosa* samples were not able to be analyzed due to DNA extractions yielding largely coral host DNA rather than microbial DNA and so are not reported on further.

Samples were collected between a depth of 2.7 – 3.9 meters. Samples were collected using a 2 cm hole-punch hammered through the live tissue. For corals that had been previously treated, samples were taken within 5 cm of the tissue margin which had previously been treated for SCTLD (Figure 1d-e). All cores were taken on angled slopes of the colonies rather than the tops or vertical sides. The hole-punch was cleaned between colonies using a disinfecting wipe. Each extracted core sample was placed into a sterile plastic bag. Upon reaching the surface, seawater was poured out of each bag and replaced with RNAlater. Bags were sealed and kept on ice back to land, then placed in a refrigerator for 24 hours to allow fixative to infiltrate the tissue, then placed in a −20°C freezer until shipment. Preserved samples were shipped frozen with cold packs to the U.S. Geological Survey’s Coral Microbial Ecology laboratory, arriving November 23, 2022, and stored in a −20°C freezer until analysis.

### 2.2. Processing and Analyses: Microbial RNAseq and 16S rRNA amplicons (Hen and Chickens Reef)

#### 2.2.1. Co-extraction of nucleic acids

Mucus-tissue slurries were homogenized using 0.1- and 0.5-mm Bashing Beads lysis tubes (Zymo Research) and vortexed horizontally for 5 min at 2500 rpm on a multi-tube vortex. DNA and RNA were co-extracted from the homogenized slurry with a ZymoBIOMICS DNA/RNA Miniprep Kit (Zymo Research) following the manufacturer instructions.

#### 2.2.2. Processing of RNAseq libraries

After extraction, total RNA and DNA were quantified spectrophotometrically using the DS-11 spectrophotometer-fluorometer tool (DeNovix). RNA integrity numbers were determined before cDNA library construction using Tape Station (Agilent). Barcoded metatranscriptomic libraries were prepared using Universal Prokaryotic RNA-seq, Any-Deplete kit (NuGen) and sequenced on two lanes of the Illumina NovaSeq6000 SP-PE-150-bp flow cell at the University of Florida’s Interdisciplinary Center for Biotechnology Research (RRID:SCR_019152). The quality of raw and filtered sequencing reads was assessed with FastQC v. 0.11.5 (Andrews 2017). Illumina adapters and quality trimming were completed using Trimmomatic v. 0.36 (Bolger et al. 2014) with the following settings: 1) a sliding window of 4 nucleotides averaging to a Phred score of 20, 2) leading and trailing sequence thresholds set to 3 nucleotides, and 3) a minimum of 50 bp in length. Ribosomal RNA sequences were removed from trimmed sequencing reads using SortMeRNA v. 4.0 (Kopylova et al. 2012). Non-ribosomal RNA sequence reads were then co-assembled with Megahit v. 1.1.4 (Li et al. 2016). Protein-coding sequences for the assembled transcripts were predicted with Prodigal v. 2.6.3 (Hyatt et al. 2010). The translated protein-coding sequences were queried against NCBI’s non-redundant protein database (downloaded in November 2022) using the Diamond blastp tool v. 2.0.14 (Buchfink et al. 2015). The top Diamond matches (highest bit score) were used for taxonomic classification with TaxonKit v. 0.9.0 (Shen and Xiong 2019). Proteins from metatranscriptomes were grouped into six taxonomic groups (Bacteria, Viruses, Archaea, Cnidaria, Symbiodiniaceae, other Eukaryotes). Antimicrobial resistance genes in proteins assigned to Bacteria were identified with RGI v. 6.0.2 with the Comprehensive Antibiotic Resistance Database (Alcock et al. 2023). Mapping to all contigs was done using Bowtie2 v. 2.4.5 (Langmead and Salzberg 2012) mapper and the transcript quantification (TPMs – transcript per million) was retrieved using Salmon v. 1.8.0 (Patro et al. 2017). The transcript quantification table was filtered to include only bacterial transcripts, and differential gene expression analysis was determined between untreated and treated samples using DESeq2 v. 1.42.1 (Love et al. 2014). The R script for analysis and plotting of the differential gene expression is available at https://github.com/meyermicrobiolab/AMR_deseq.

#### 2.2.3 Processing of 16S rRNA amplicon libraries

The bacterial community composition of both RNA and DNA fractions of each sample was characterized by amplicon sequencing of the V4 region of the 16S rRNA gene. Extracted RNA was converted to cDNA with a ProtoScript II First strand cDNA synthesis kit (New England Biolabs) before use as a template for PCR. Amplicon libraries were generated as previously described (Meyer et al. 2019) using the 515F (Parada et al. 2016) and 806RB (Apprill et al. 2015) Earth Microbiome primers. Amplification was performed in triplicate 25-μl reactions for each sample with Phusion High-Fidelity Master Mix (New England Biolabs) with 3% dimethyl sulfoxide, 0.25μM of each primer, and 2μl template. Triplicate reactions were cleaned with a Qiagen MinElute PCR purification kit and quantified with a DeNovix DS-11 spectrophotometer. One amplicon pool was generated with 240 ng of each cleaned, barcoded PCR product for sequencing at the University of Florida’s Interdisciplinary Center for Biotechnology Research (RRID:SCR_019152) on an Illumina MiSeq with 2 × 150bp v. 2 cycle format. Illumina adapters and primers were removed from raw sequencing reads with cutadapt v. 1.8.1 (Martin 2011) and remaining analyses were completed in RStudio v. 2023.12.1+402 with R version 4.3.2. Amplicon sequence variants were determined with DADA2 v. 1.30.0 (Callahan et al. 2016). Taxonomy was assigned with the Silva nr99 v. 138.1 database (Yilmaz et al. 2013). Low abundance amplicon sequence variants with a mean read count across all samples of less than five were removed from the analysis. Sequencing read count data were transformed by center-log-ratio (CLR) and the Aitchison distance of CLR-transformed data was determined with CoDaSeq (Gloor et al. 2017) and microbial communities were analyzed with phyloseq v. 1.46.0 (McMurdie and Holmes 2013) and vegan v. 2.6-4 (Dixon 2003). The full reproducible R script is available at https://github.com/meyermicrobiolab/AMR-16S-rRNA.

### 2.3 Processing and Analyses: Microbial qPCR Array (Cheeca Rocks)

Using sterile technique, preserved coral biopsies were thawed and crushed using a sterile hammer, and approximately 0.2g of each sample was transferred to a bead-beating tube. DNA extractions were conducted using Qiagen DNeasy PowerBiofilm kits following the manufacturer’s protocol. This methodology has been previously used in similar assessments in mariculture (Zhao et al. 2017) and urban rivers (Yitayew et al. 2022). DNA concentrations were quantified using the dsDNA Broad Range Qubit 3.0 fluorometric assay. Samples that yielded DNA concentrations below 10 ng/µL were re-extracted and quantified.

DNA samples were screened for antibiotic resistance genes using Qiagen Microbial DNA qPCR array for Antibiotic Resistance Genes (ARGs). The assays in each of the 96-wells of the microarray contain predispensed PCR primers and 5’-hydrolysis probe sets that enable quantitative real-time PCR. Each microarray plate contains assays for 87 antibiotic resistance genes belonging to aminoglycoside, beta-lactam, erythromycin, fluoroquinolone, macrolide-lincosamide-streptogramin B, tetracycline, vancomycin, and multidrug resistance classifications (Table S1). Beta-lactam resistance genes protect bacteria by hydrolyzing the beta-lactam ring of various compounds, including amoxicillin. The array contains reactions for 57 beta-lactam resistance genes of different classes based on functions and sequence similarity. Additionally, each plate includes positive control wells for the presence of bacterial DNA and positive PCR controls to test for the presence of inhibitors and confirm the efficiency of the reaction. For each array, a reaction mixture was made consisting of 1,275 µL of Qiagen Microbial qPCR Mastermix, 94 µL of sample DNA, and 1,181 µL of Microbial DNA-Free Water, for a total volume of 2,550 µL which allows 25 µL of reaction mix to be added to each well using a multichannel pipet and reagent reservoir. The manufacturer recommends using a minimum of 500 ng genomic DNA from metagenomic samples.

Given that it is not possible to determine how much of the sample DNA concentration was coral host DNA versus bacterial microbiome DNA, we applied the maximum concentration available from each sample (range: 1,100–10,246 ng/µL; median: 2,634 ng/µL). The microarrays were run on an Applied Biosystems StepOnePlus running software version 2.2.2, with baseline set as 8–20 cycles and a threshold setting of 0.2. The cycling conditions consisted of a single initial PCR activation step of 95°C for 10 min, followed by 40 cycles of 95°C for 15 sec denaturation and 60°C for 2 min annealing and extension. Raw C_t_ values were exported to the Microbial qPCR analysis Excel template to detect the presence of ARGs. The identification criteria were C_T_ <32 = positive, C_t_ >37 = negative, and C_t_ >32–<37 = inconclusive (Zhao et al. 2017). Quality controls included that the six PanBacteria wells had C_t_ <37 (confirming there was sufficient bacterial DNA in the sample to detect ARGs if present in sufficient gene copy numbers) and the positive PCR control (PPC) wells needed a C_t_ = 22±2 to confirm the instrument and mastermix performed correctly and that there were no PCR inhibitors in the sample. Per the manufacturer’s documentation, most of the ARGs assays had a sensitivity of 20–50 gene copies per target, with a small number of targets requiring 100–200 gene copies for detection (Table S1). Exact sample DNA concentration values, raw fluorescence values, and raw threshold cycle (C_t_) values are available for each sample in the U.S. Geological Survey data release (Kellogg and Voelschow 2024).

## 3. Results

### 3.1 Microbial RNAseq and 16S rRNA amplicons (Hen and Chickens Reef)

Nucleic acids were extracted from a total of twenty mucus-slurries that were collected from ten *M. cavernosa* colonies before and 24 hours after the application of an antibiotic treatment. RNAseq libraries were successfully generated from 13 out of the 20 samples with RNA integrity numbers > 0.8. A total of 514,841,463 sequencing reads were generated for the 13 RNAseq libraries, with an average of 34,838,281 reads per sample after quality-filtering. Assembled RNAseq libraries including genetic sequences with gene annotations are publicly available and searchable in the Joint Genome Institute’s Integrated Microbial Genomes and Microbiomes database under the accession numbers listed in Table S2. Amplicon libraries were successfully generated from 19 out of 20 DNA fractions and 18 out of 20 RNA fractions. A total of 1,428,607 sequencing reads were generated for the 37 amplicon libraries, with an average of 15,409 reads per sample after quality-filtering. All raw sequencing reads are available in NCBI under Bioproject PRNJA931253.

A total of 42,770 bacterial genes encoding antimicrobial resistance mechanisms were expressed. The most commonly expressed resistance mechanisms were antibiotic efflux and antibiotic inactivation (Figure 2). A total of 12,824 transcripts for antibiotic efflux genes were detected, which are used to actively pump out a broad range of antimicrobial compounds. A total of 17,901 transcripts for antibiotic inactivation genes were detected, which act upon specific classes of antimicrobial compounds. The third most commonly expressed resistance mechanism was antibiotic target alteration. Of the 6,726 transcripts annotated as antibiotic target alteration, 57 transcripts were annotated as penicillin-binding protein mutations conferring resistance to beta-lactam antibiotics. A total of 261 gene families for antibiotic inactivation were expressed, of which 202 gene families were beta-lactamases that can inactivate beta-lactam antibiotics like amoxicillin. Expressed antimicrobial inactivation gene families were very diverse and the most common included OXA beta-lactamase and nitroimidazole reductase (Figure 3).

**Figure 2.**
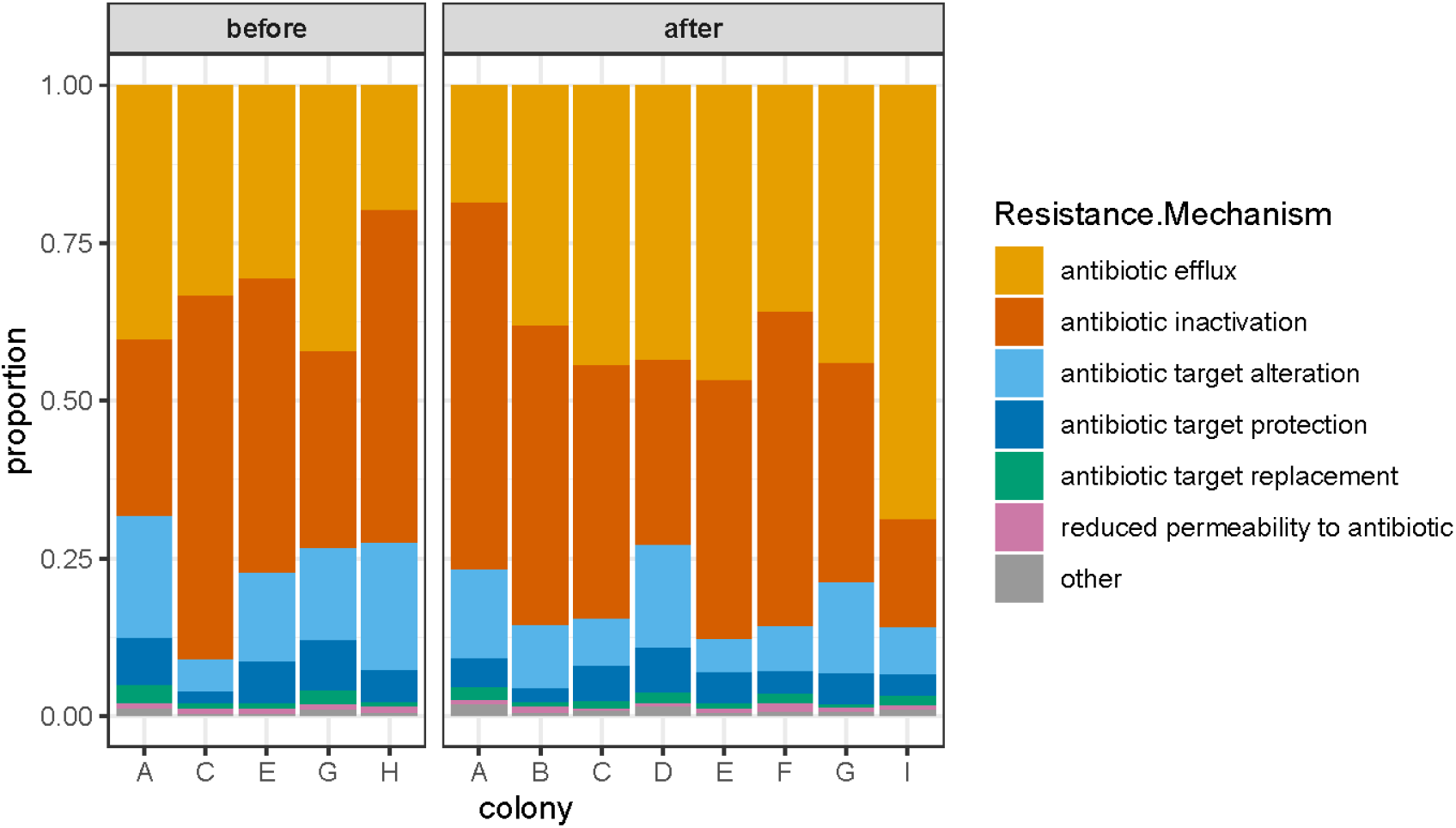
Proportion of expressed antimicrobial resistance genes per resistance mechanism for *Montastraea cavernosa* colonies before and 24 hours after treatment.

**Figure 3.**
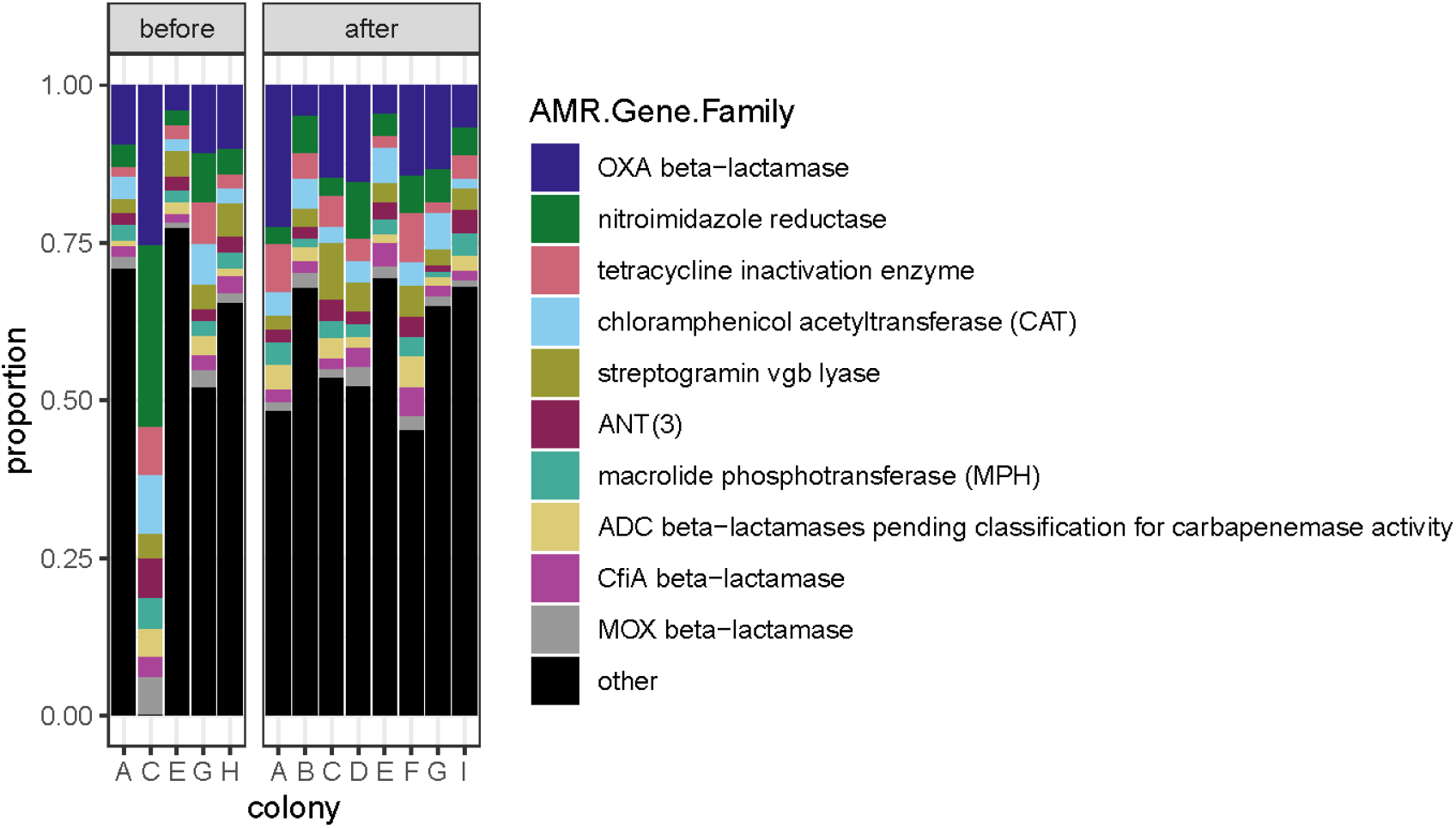
Proportion of expressed antimicrobial inactivation gene families in *Montastraea cavernosa* colonies before and 24 hours after treatment. The ten most abundant gene families by gene count are shown; the remaining 203 antimicrobial inactivation gene families are included as “other”.

To determine what background ARG levels were, and if the amoxicillin treatments induced short-term changes in the expression of ARGs, we tested differential expression between samples collected before the treatment and those collected one day later. No differentially expressed ARGs were detected. Furthermore, when ARGs were grouped by drug class, resistance mechanism, or resistance gene family, no differentially expressed classes, mechanisms, or families were detected.

In addition to determining if ARG expression was impacted by amoxicillin treatment, we also examined the microbiome composition based on both DNA and RNA fractions before and after amoxicillin treatment. Overall, 11% of the variation in the microbial community structure was explained by the nucleic acid fraction (PERMANOVA R^2^ = 0.11, p = 0.002) while only 7% of the variation was explained by when the sample was taken (before or after treatment) (PERMANOVA R^2^ = 0.07, p = 0.018) (Figure 4). Thus, 82% of the community variation was not explained by either factor.

**Figure 4.**
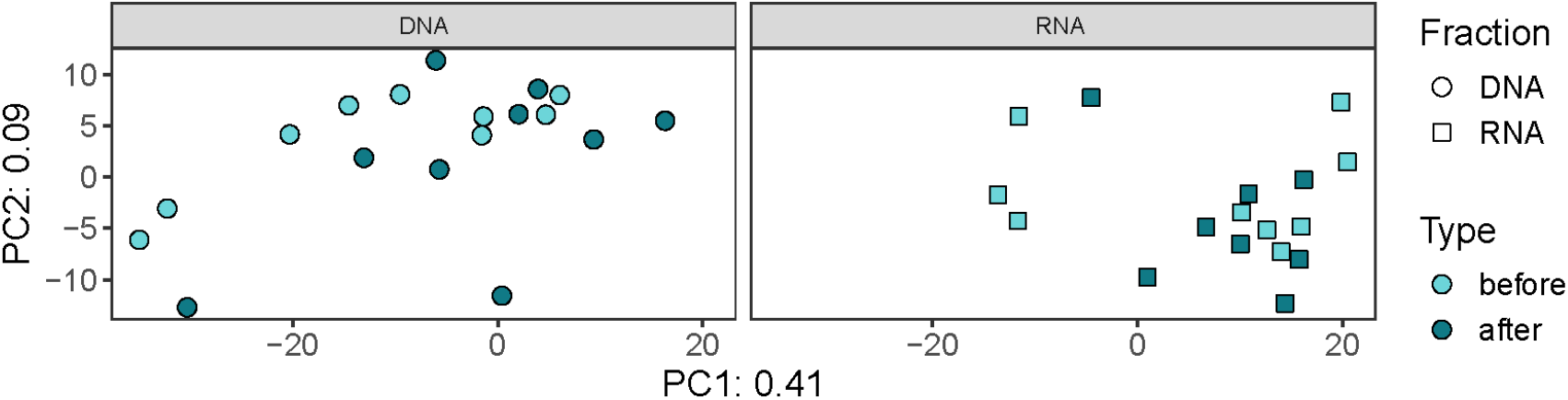
Principal component analysis of microbial communities in *Montastraea cavernosa* corals before and after amoxicillin treatment, faceted by nucleic acid fraction. Half of the variation among communities is shown by the first two principal components (PC1 and PC2).

In both the RNA and DNA fractions, *Montastraea cavernosa* microbial communities at Hen and Chickens reef were composed primarily of Alphaproteobacteria and Gammaproteobacteria (Figure S1). Microbial communities in the DNA fraction were composed predominantly of members of the SAR11 clade, Flavobacteriales, and Blastocatellales, while the RNA fraction was predominantly composed of Pseudomonadales, Sphingomonadales, and an unclassified order of Proteobacteria (Figure 5).

**Figure 5.**
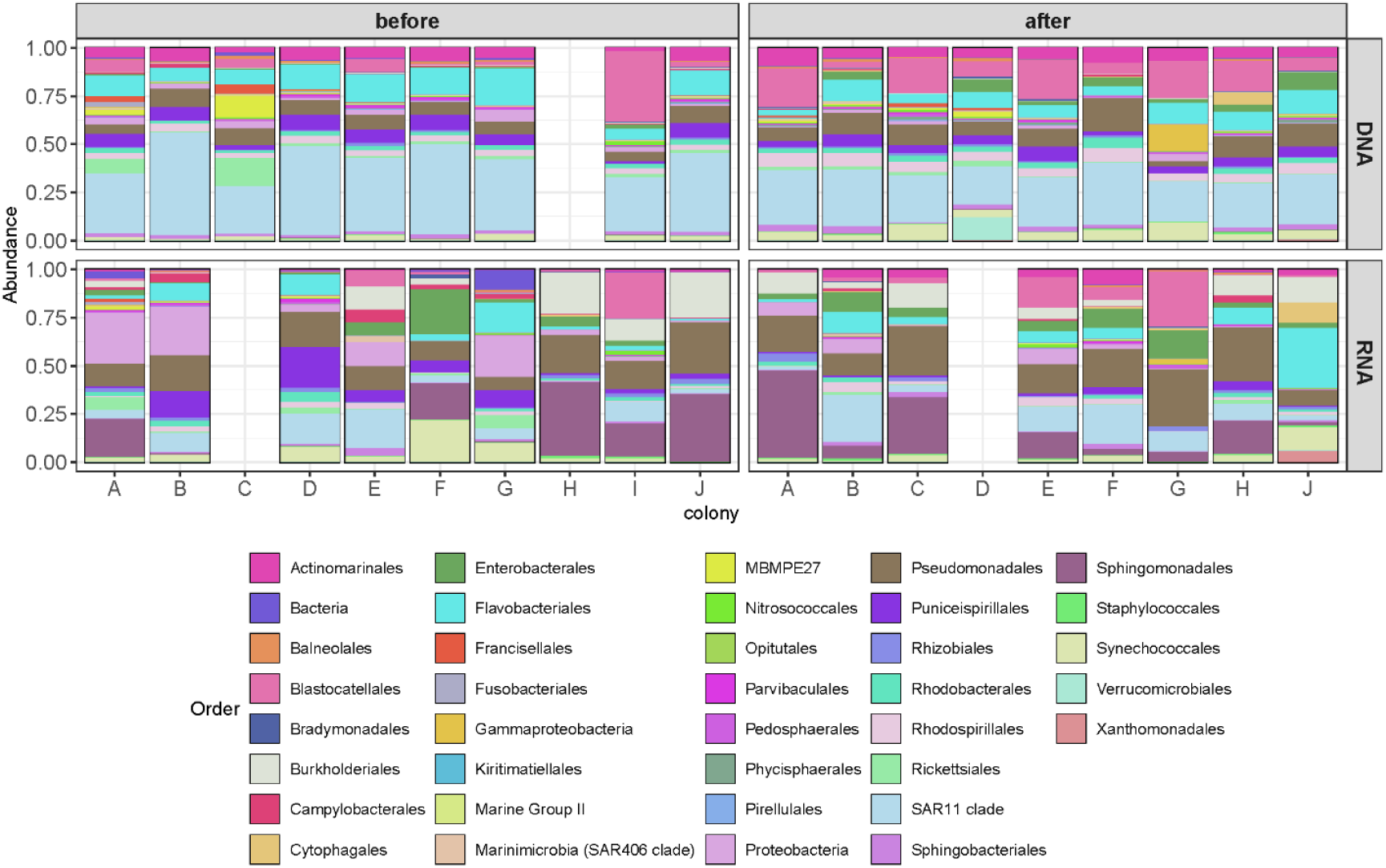
Relative abundance of amplicon sequence variants, colored by Order or lowest taxonomic classification possible, in *Montastraea cavernosa* samples before and after amoxicillin treatment. Community composition of the DNA fraction is shown on top while composition of the RNA fraction is shown below for each colony (designated by capital letters).

### 3.2 Microbial qPCR Array (Cheeca Rocks)

All array plates that were run met the quality requirement that the three positive PCR control (PPC) replicate assays reached a C_t_ = 22±2 to confirm the instrument and mastermix performed correctly and that there were no PCR inhibitors in the samples. However, the *M. cavernosa* DNA samples did not meet the criteria of C_t_ <37 for the triplicate assays for the PanBacteria 1 and PanBacteria 3 controls and therefore the *M. cavernosa* data were not considered valid. Note that this is likely due to *M. cavernosa* having large polyps which generate a lot of host DNA that can swamp the microbial DNA signal. Conversely, the *C. natans* DNA samples did meet the criteria for C_t_ <37 in the triplicate assays for the PanBacteria 1 and PanBacteria 3 controls, confirming there was sufficient bacterial DNA in the samples for ARGs detection: PanBacteria1 wells had a range of 24.18–31.6, with a median value of 28.41 and PanBacteria 3 wells had a range of 25.58–33.38, with a median value of 30.49. Based on the stated identification criteria of C_t_ <32 = positive, C_t_ >37 = negative, and C_t_ >32–<37 = inconclusive, there were zero ARGs detected in the 12 *C. natans* samples. However, there were a small number of above-threshold detections of three genes, *ermB* (20 copy sensitivity), *mefA* (100 copy sensitivity), and *msrA* (100 copy sensitivity) all of which belong to the classification macrolide-lincosamide-streptogramin_b (MLS) (Figure 6). Since these assays had extremely high C_t_ values (38.32–39.51) they may have resulted from non-specific amplification. It is, however, interesting to note that of the six times they occurred, four were in samples from corals that had never been treated, and the remaining two were in corals that had been treated once; none occurred in corals that had been treated multiple times.

**Figure 6.**
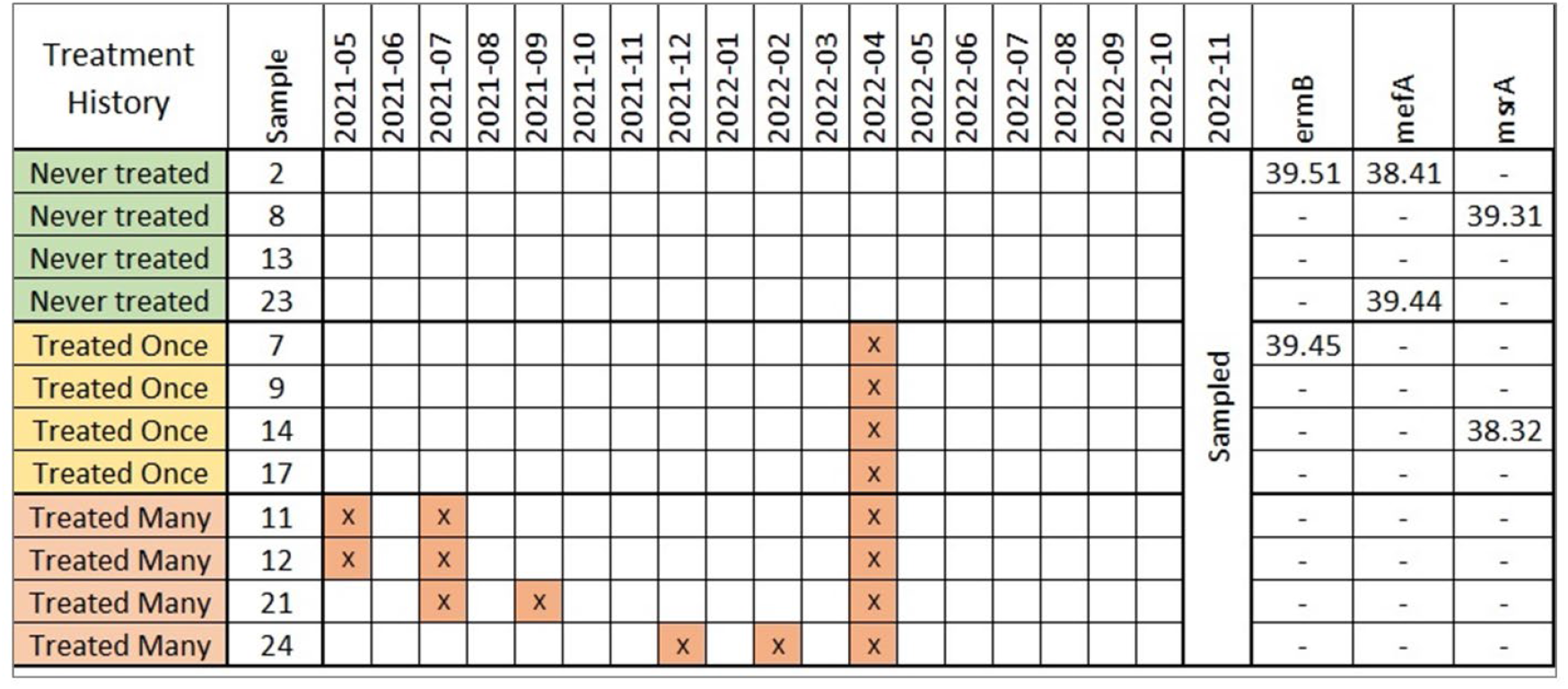
Treatment history and above-threshold ARGs detections for *Colpophyllia natans* coral colonies sampled for microbial qPCR array. Shaded boxes with an x indicate date (year-month) when a coral was treated prior to November 2022 sampling. Genes observed outside the acceptable identification criteria (C_t_ >37) are listed with their C_t_ values.

## 4. Discussion

These two independently designed experiments were intended to assess the real-world impacts of topical antibiotic applications on ARGs in wild SCTLD-affected corals. We specifically targeted two distinct time points - 24 hours after application and seven months after application - to address both immediate and long-term potential for the development of antibiotic resistance and impacts of intervention efforts on the coral microbiome.

There are several common analytical methods used to investigate ARGs. In clinical settings it is common to first isolate candidate microbes and then screen them for resistance to specific antibiotics. While this technique can be applied to coral-associated bacteria (e.g., Galkiewicz et al. (2011)), it is limited to those bacterial taxa that are amenable to cultivation. For environmental assessments, molecular techniques including targeted quantitative polymerase chain reaction (qPCR) and metagenomics are the preferred methods (Scott et al. 2020, Davis et al. 2023). While molecular techniques cannot assess bacterial viability, they can be scaled to enable high-throughput of larger sample sizes. Targeted genomic analysis via the commercial qPCR microarray used in this study allows for rapid screening of more than 80 clinically relevant ARGs at detection limits ranging from 20–100 gene copies (Table S1). Running a sample takes approximately two hours, followed by minimal analysis using a spreadsheet. However, this system relies on having prior sequence knowledge of the target ARGs and can only detect the specifically chosen ARGs on the array, thus limiting the scope of detection to these clinically relevant ARGs. In contrast, metagenomic analysis is an open-ended tool that can reveal previously undiscovered diversity of ARGs in an environmental microbial community. However, the post-sequencing bioinformatic analysis is much more time- and computation-intensive. In addition, metagenomic studies examining bacterial genes remain technically challenging to conduct with coral microbiota because of the overwhelming genetic signature of the host (Messer et al. 2024). Therefore, we used a metatranscriptomic approach which more effectively targets the bacterial RNA fraction and provides a snapshot of actively transcribed genes. While this approach revealed the highly diverse ARG repertoire of coral microbes, successful recovery of high-quality RNA libraries was challenging and resulted in lowered statistical power. Therefore, we recommend that future efforts include multiple sample replicates from the same colony if permitting allows.

The rationale for using a microarray with 87 target ARGs, of which 57 are beta-lactamase genes (relevant to amoxicillin; inclusive of classes A, B, C, and D) was to determine if: 1) there was an increase in abundance or diversity of ARGs directly linked to amoxicillin resistance and 2) whether, due to multiple ARGs being carried on plasmids, this led to an increase in ARGs to other classes of antibiotics. Instead, there were zero verified detections for clinical ARGs based on microarrays. We did observe late amplification (past the cut off criteria of C_t_ < 37; Figure 6) in five samples for genes (*ermB, mefA, msrA*) within the macrolide-lincosamide-streptogramin (MLS) group of resistance genes. These only occurred in the never-treated or treated-once corals. This might be non-specific amplification, but it is worth noting that this group of genes is extremely common in environmental bacteria (defined as genera and species primarily found outside of humans and animals) (Roberts 2011). We surmise that these late amplifications could be instances of array primers picking up environmental genes that are similar but not exact matches to their optimized targets (Aminov 2009). Persistence of certain bacteria during laboratory experiments involving antibiotic treatment of corals implied the presence of some naturally antibiotic-resistant bacteria in coral microbiomes (Sweet et al. 2011, Connelly et al. 2022, Connelly et al. 2023).

To date, only a handful of studies have examined ARGs associated with corals or coral reef ecosystems, but these all suggest that the presence of ARGs in marine bacteria is likely common and widespread. For example, isolates of the coral pathogen *Vibrio coralliilyticus* have demonstrated antimicrobial resistance to most of the 26 clinically relevant antibiotics, including amoxicillin (Vizcaino et al. 2010). In addition, ARGs for eight clinically relevant antibiotics were detected by quantitative PCR in microorganisms from both seawater (Liu et al. 2020) and microplastics (Zhou et al. 2024) from coral reefs in China. Here, even with fewer than 20 metatranscriptomic libraries, we detected the expression of over 42,000 unique bacterial ARGs. The most widespread resistance to beta-lactams is through beta-lactamases (Williams 1999), which were discovered in 1940 (Abraham and Chain 1940) as soon as penicillin was introduced to clinical practice. Given the widespread and well-studied nature of beta-lactamases, it is not surprising that we found genes for resistance to amoxicillin, even in untreated corals. These findings suggest that expressed ARGs are extremely diverse and common in the environment, regardless of small-scale, localized treatments. Furthermore, this study provides an extensive dataset of these ARGs that may serve as a resource for future studies. Specifically, the untargeted RNAseq approach provided the genetic sequences of ARGs that are present in Florida corals which can be used to expand the scope of detection in future qPCR assays.

Overuse and misuse of antibiotics in human and veterinary medicine, agriculture, and aquaculture are considered the main cause of the rise in antimicrobial-resistant pathogens and development of environmental reservoirs of clinically relevant ARGs (Thanner et al. 2016, Dadgostar 2019). The contributions of topical antimicrobials to this rise are largely unknown due to insufficient data (Blackburn et al. 2023). Beta-lactam antibiotics like amoxicillin make up the largest share of human-use antibiotics in most countries, accounting for more than half of total antibiotic use (Kümmerer 2009a). Beta-lactam ARGs are the most commonly detected resistance genes in the marine environment, based on a meta-analysis of metagenomic data from different biomes (Zhang et al. 2022). However, high amounts of clinically relevant ARGs are primarily detected in areas of high human interaction, such as hospitals, wastewater, and areas directly impacted by large scale farming (Zhang et al. 2022). The amount of selective pressure being exerted by a spatially limited dose of antibiotic paste, compared to these sources, or to an aquaculture situation where higher doses of antibiotics are being consistently employed either in medicated feed or via bath immersion (Chen et al. 2018), is a comparatively negligible addition to the environment.

The reflexive expectation that treating corals with antibiotic paste would result in selective pressure favoring antibiotic-resistant bacteria and promotion of horizontal gene transfer of ARGs via plasmids needs to be examined. We show here that the small, focused treatments used to treat SCTLD-affected corals are not increasing the diversity or abundance of ARGs at either immediate (24 hour) nor longer-term (7+ month) time frames. One potential consideration for the absence of change to the microbiome or increase in ARGs in the samples taken 24 hours after treatment is that the 2 cm distance between the treatment and the sample was too far for effects to be seen. However, we think this is unlikely based on the interconnectedness of coral polyps. Though distinct from scleractinian corals, Gateno et al. (1998) found material translocation through the gastrovascular system of octocorals across 4 cm of tissue in 24 hours. Neely and Lewis (2018) also observed a progressing SCTLD lesion on the scleractinian coral *Dendrogyra cylindrus* halting 2-5 cm from an amoxicillin-packed trench, suggesting effective dosages reached that distance from the treatment line.

As elevated ARGs are not found even in the treatment-adjacent coral tissues, it is extremely unlikely that they could be dispersing farther afield into surrounding organisms, sediment, or the water column. Beta-lactams are typically only detected in low concentrations in the environment and are easily hydrolyzed at ambient temperature (Längin et al. 2009, Bergheim et al. 2010). In addition, the unstable beta-lactam ring is inactivated by both heat and UV radiation (Kovalakova et al. 2020). This suggests that any amoxicillin that leaches out of the amoxicillin+Base2b paste is not only rapidly diluted by seawater but also chemically broken down.

There is evidence from a variety of studies that the coral-associated microbial community can be resilient to environmental stressors and can return to the original community composition after disturbance (Garren et al. 2009, Ziegler et al. 2019, Strudwick et al. 2022). After deliberate removal of more than 99% of the coral-associated bacteria via a 6-day immersion treatment of ciprofloxacin, Sweet et al. (2011) identified substantial recovery of the bacterial community by day 4 (the end of the experimental monitoring). In another experiment that treated corals in aquaria with an antibiotic mixture for 8 days and then returned them to the reef, most of the treated corals succumbed to bleaching or mortality, but the few survivors regained a mucus microbiome similar to pretreatment by 28 days (Glasl et al. 2016). A similar experiment that exposed corals to an antibiotic cocktail for a week found that the mucus microbiome recovered to its initial state within 2 weeks (Bent et al. 2021). Our results suggest a negligible effect of treatments on the microbiome 24 hours after treatment, probably due to the topical nature of application. The use of a slow-release antibiotic paste applied only to lesions, rather than full colony immersion in liquid antibiotics for 3 to 7 days, did not result in the dramatic reduction in bacterial diversity seen in prior laboratory experiments (Sweet et al. 2011, Gilbert et al. 2012, Sweet et al. 2014, Glasl et al. 2016, Connelly et al. 2022, Connelly et al. 2023). The effects of antibiotics on coral health via disruption of the microbiome observed in prior tank experiments appear to be a direct consequence of both the longer antibiotic exposure times and the choice to uniformly expose the entire coral fragment.

SCTLD is known to be extremely lethal to affected corals (Precht et al. 2016, Aeby et al. 2019, Thome et al. 2021), and in-water topical antibiotic intervention offers the only known tool for preventing that mortality. As managers are considering the use of this tool and weighing the pros with the potential cons of its use, the spread of antibiotic resistance may be a topic of discussion. This study identifies that within the parameters (location, species, and timeframes) we tested, development of antibiotic resistance through these treatments is unlikely to be a factor of concern.

## 5. Acknowledgments

We are grateful for additional field assistance by Sydney Gallagher and Michelle Dobler. This work was supported by the U.S. Geological Survey’s Ecosystems Biological Threats and Invasive Species Research Program (CAK). Funding support was also provided by Florida Department of Environmental Protection Office of Resilience and Coastal Protection – Southeast Region (JM). Collections were conducted opportunistically under a Florida DEP grant (KLN). Additional time was donated by the author (KLN). All work was conducted under Florida Keys National Marine Sanctuary permits FKNMS-2022-066-A1 and FKNMS-2019-177-A3. Any use of trade, firm, or product names is for descriptive purposes only and does not imply endorsement by the U.S. Government.

## 6. Data Availability Statement

The raw data supporting the conclusions of this article will be made available by the authors, without undue reservation. Raw sequencing reads from the RNAseq and 16S rRNA amplicon studies are publicly available at NCBI under BioProject PRJNA931253. Assembled RNAseq libraries including genetic sequences with gene annotations are publicly available and searchable in the Joint Genome Institute’s Integrated Microbial Genomes and Microbiomes database. The R scripts for analysis and plotting of the differential gene expression is available at https://github.com/meyermicrobiolab/AMR_deseq. The R scripts for analysis of microbiome composition is available at https://github.com/meyermicrobiolab/AMR-16S-rRNA. Raw fluorescence data and cycle threshold data from microarray study is publicly available via USGS data release https://doi.org/10.5066/P9AST28V.

## 8. Figures

**Figure S1.**
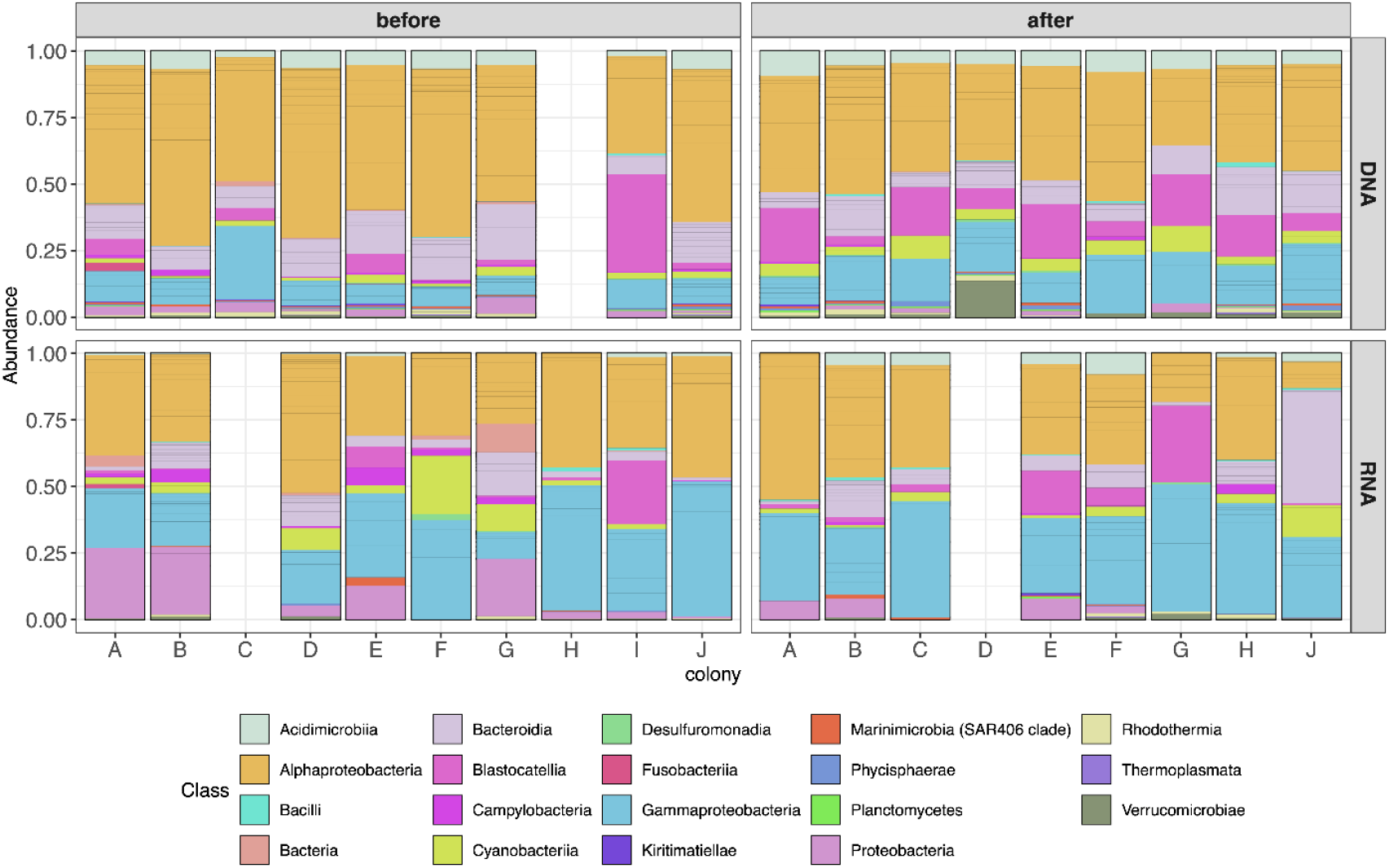
Relative abundance of amplicon sequence variants, colored by Class, in Montastraea *cavernosa* corals before and 24 hours after amoxicillin treatment. Community composition of the DNA fraction is shown on top while composition of the RNA fraction is shown below for each colony.

**Table S1.** List of assays in the microbial qPCR array. Well is the position in the 96-well plate, Species/Gene is the target of the assay, and Sensitivity is the minimum number of gene copies required for detection. Pan Bacteria 1 and 3 are internal controls for detection of bacterial DNA and PPC is the positive PCR control that confirms no inhibition of amplification.

| Well | Species/Gene | Antibiotic classification | Sensitivity |
| --- | --- | --- | --- |
| A1 | AAC(6)-Ib-cr | Fluoroquinolone resistance | 100 |
| A2 | aacC1 | Aminoglycoside resistance | 50 |
| A3 | aacC2 | Aminoglycoside resistance | 30 |
| A4 | aacC4 | Aminoglycoside resistance | 20 |
| A5 | aadA1 | Aminoglycoside resistance | 200 |
| A6 | aphA6 | Aminoglycoside resistance | 40 |
| A7 | BES-1 | Class A beta-lactamase | 20 |
| A8 | BIC-1 | Class A beta-lactamase | 100 |
| A9 | CTX-M-1 Group | Class A beta-lactamase | 50 |
| A10 | CTX-M-8 Group | Class A beta-lactamase | 40 |
| A11 | CTX-M-9 Group | Class A beta-lactamase | 30 |
| A12 | GES | Class A beta-lactamase | 20 |
| B1 | IMI & NMC-A | Class A beta-lactamase | 30 |
| B2 | KPC | Class A beta-lactamase | 40 |
| B3 | Per-1 group | Class A beta-lactamase | 30 |
| B4 | Per-2 group | Class A beta-lactamase | 50 |
| B5 | SFC-1 | Class A beta-lactamase | 50 |
| B6 | SFO-1 | Class A beta-lactamase | 20 |
| B7 | SHV | Class A beta-lactamase | 200 |
| B8 | SHV(156D) | Class A beta-lactamase | 100 |
| B9 | SHV(156G) | Class A beta-lactamase | 50 |
| B10 | SHV(238G240E) | Class A beta-lactamase | 30 |
| B11 | SHV(238G240K) | Class A beta-lactamase | 40 |
| B12 | SHV(238S240E) | Class A beta-lactamase | 50 |
| C1 | SHV(238S240K) | Class A beta-lactamase | 30 |
| C2 | SME | Class A beta-lactamase | 30 |
| C3 | TLA-1 | Class A beta-lactamase | 50 |
| C4 | VEB | Class A beta-lactamase | 20 |
| C5 | ccrA | Class B beta-lactamase | 30 |
| C6 | IMP-1 group | Class B beta-lactamase | 50 |
| C7 | IMP-12 group | Class B beta-lactamase | 200 |
| C8 | IMP-2 group | Class B beta-lactamase | 30 |
| C9 | IMP-5 group | Class B beta-lactamase | 200 |
| C10 | NDM | Class B beta-lactamase | 50 |
| C11 | VIM-1 group | Class B beta-lactamase | 50 |
| C12 | VIM-13 | Class B beta-lactamase | 20 |
| D1 | VIM-7 | Class B beta-lactamase | 40 |
| D2 | ACC-1 group | Class C beta-lactamase | 100 |
| D3 | ACC-3 | Class C beta-lactamase | 30 |
| D4 | ACT 5/7 group | Class C beta-lactamase | 30 |
| D5 | ACT-1 group | Class C beta-lactamase | 100 |
| D6 | CFE-1 | Class C beta-lactamase | 50 |
| D7 | CMY-10 Group | Class C beta-lactamase | 30 |
| D8 | DHA | Class C beta-lactamase | 20 |
| D9 | FOX | Class C beta-lactamase | 100 |
| D10 | LAT | Class C beta-lactamase | 100 |
| D11 | MIR | Class C beta-lactamase | 30 |
| D12 | MOX | Class C beta-lactamase | 30 |
| E1 | OXA-10 Group | Class D beta-lactamase | 20 |
| E2 | OXA-18 | Class D beta-lactamase | 30 |
| E3 | OXA-2 Group | Class D beta-lactamase | 40 |
| E4 | OXA-23 Group | Class D beta-lactamase | 50 |
| E5 | OXA-24 Group | Class D beta-lactamase | 20 |
| E6 | OXA-45 | Class D beta-lactamase | 100 |
| E7 | OXA-48 Group | Class D beta-lactamase | 50 |
| E8 | OXA-50 Group | Class D beta-lactamase | 20 |
| E9 | OXA-51 Group | Class D beta-lactamase | 100 |
| E10 | OXA-54 | Class D beta-lactamase | 20 |
| E11 | OXA-55 | Class D beta-lactamase | 30 |
| E12 | OXA-58 Group | Class D beta-lactamase | 20 |
| F1 | OXA-60 | Class D beta-lactamase | 30 |
| F2 | ereB | Erythromycin resistance | 20 |
| F3 | QepA | Fluoroquinolone resistance | 50 |
| F4 | QnrA | Fluoroquinolone resistance | 40 |
| F5 | QnrB-1 group | Fluoroquinolone resistance | 20 |
| F6 | QnrB-31 group | Fluoroquinolone resistance | 20 |
| F7 | QnrB-4 group | Fluoroquinolone resistance | 30 |
| F8 | QnrB-5 group | Fluoroquinolone resistance | 40 |
| F9 | QnrB-8 group | Fluoroquinolone resistance | 20 |
| F10 | QnrC | Fluoroquinolone resistance | 30 |
| F11 | QnrD | Fluoroquinolone resistance | 40 |
| F12 | QnrS | Fluoroquinolone resistance | 40 |
| G1 | ermA | Macrolide Lincosamide Streptogramin_b | 100 |
| G2 | ermB | Macrolide Lincosamide Streptogramin_b | 20 |
| G3 | ermC | Macrolide Lincosamide Streptogramin_b | 100 |
| G4 | mefA | Macrolide Lincosamide Streptogramin_b | 100 |
| G5 | msrA | Macrolide Lincosamide Streptogramin_b | 100 |
| G6 | oprj | Multidrug resistance efflux pump | 50 |
| G7 | oprm | Multidrug resistance efflux pump | 20 |
| G8 | tetA | Tetracycline efflux pump | 40 |
| G9 | tetB | Tetracycline efflux pump | 30 |
| G10 | vanB | Vancomycin resistance | 100 |
| G11 | vanC | Vancomycin resistance | 30 |
| G12 | *Staphylococcus aureus* |  | 100 |
| H1 | mecA | Beta-lactam resistance | 40 |
| H2 | lukF | Panton Valentine leukocidin chain F precursor | 20 |
| H3 | spa | Immunoglobulin G binding protein A precursor | 200 |
| H4 | Pan Bacteria 1 | n/a | n/a |
| H5 | Pan Bacteria 1 | n/a | n/a |
| H6 | Pan Bacteria 1 | n/a | n/a |
| H7 | Pan Bacteria 3 | n/a | n/a |
| H8 | Pan Bacteria 3 | n/a | n/a |
| H9 | Pan Bacteria 3 | n/a | n/a |
| H10 | PPC | n/a | n/a |
| H11 | PPC | n/a | n/a |
| H12 | PPC | n/a | n/a |

**Table S2.**
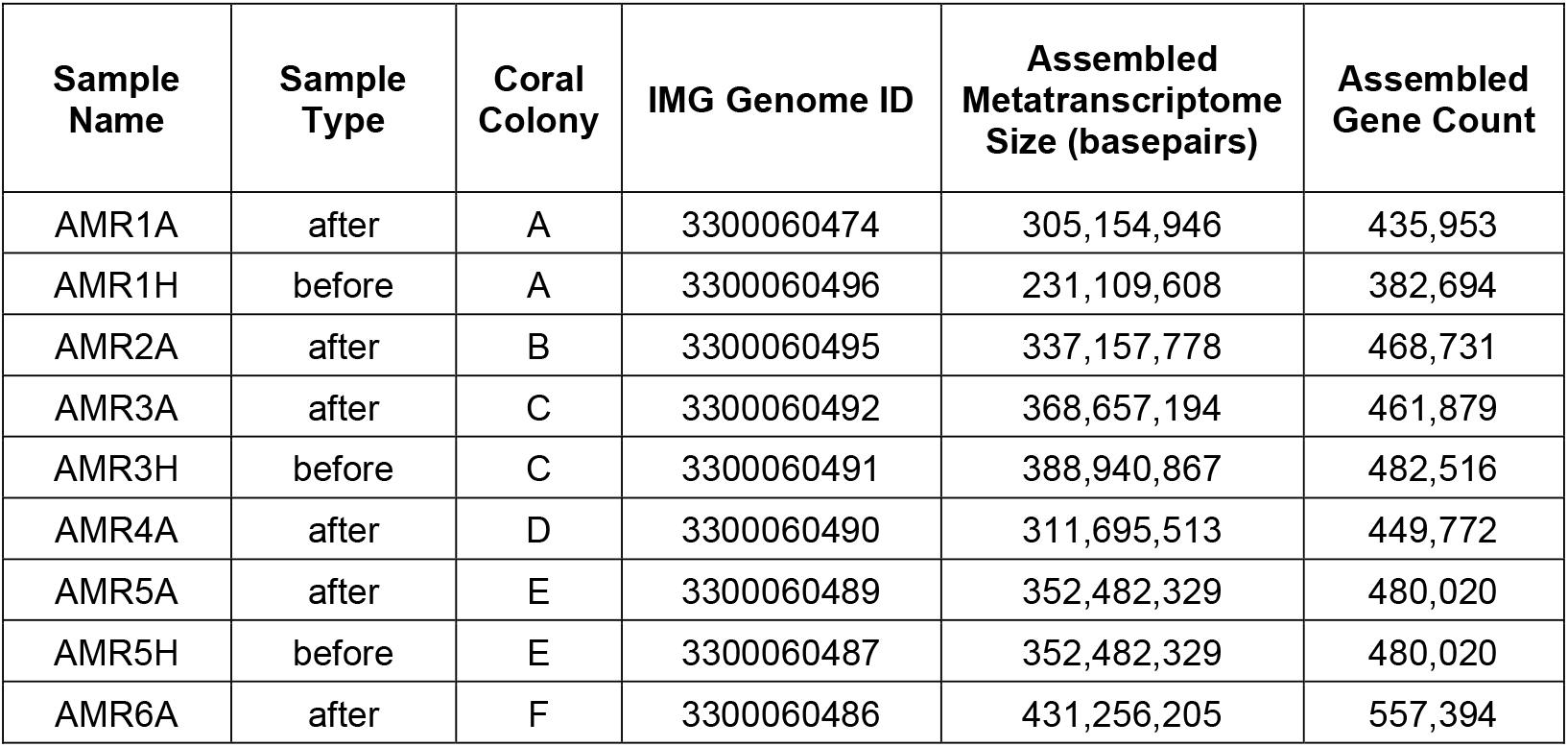
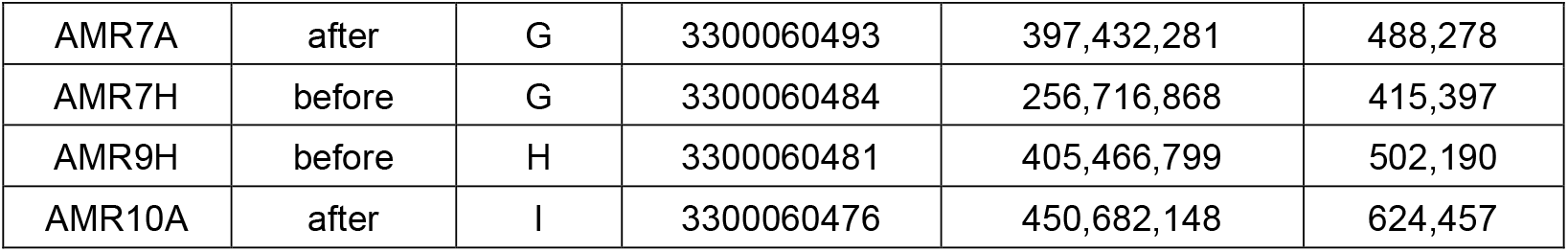
Assembled RNAseq libraries from this study that are publicly available in the Joint Genome Institute’s Integrated Microbial Genomes and Microbiomes database (img.jgi.doe.gov)

| Sample Name | Sample Type | Coral Colony | IMG Genome ID | Assembled Metatranscriptome Size (basepairs) | Assembled Gene Count |
| --- | --- | --- | --- | --- | --- |
| AMR1A | after | A | 3300060474 | 305,154,946 | 435,953 |
| AMR1H | before | A | 3300060496 | 231,109,608 | 382,694 |
| AMR2A | after | B | 3300060495 | 337,157,778 | 468,731 |
| AMR3A | after | C | 3300060492 | 368,657,194 | 461,879 |
| AMR3H | before | C | 3300060491 | 388,940,867 | 482,516 |
| AMR4A | after | D | 3300060490 | 311,695,513 | 449,772 |
| AMR5A | after | E | 3300060489 | 352,482,329 | 480,020 |
| AMR5H | before | E | 3300060487 | 352,482,329 | 480,020 |
| AMR6A | after | F | 3300060486 | 431,256,205 | 557,394 |
| AMR7A | after | G | 3300060493 | 397,432,281 | 488,278 |
| AMR7H | before | G | 3300060484 | 256,716,868 | 415,397 |
| AMR9H | before | H | 3300060481 | 405,466,799 | 502,190 |
| AMR10A | after | I | 3300060476 | 450,682,148 | 624,457 |

